# Interaction of human keratinocytes and nerve fiber terminals at the neuro-cutaneous unit

**DOI:** 10.1101/2022.02.23.481592

**Authors:** Christoph Erbacher, Sebastian Britz, Philine Dinkel, Thomas Klein, Markus Sauer, Christian Stigloher, Nurcan Üçeyler

**Affiliations:** Department of Neurology, University of Würzburg, 97080 Würzburg, Germany; Imaging Core Facility, Biocenter, University of Würzburg, 97074 Würzburg, Germany; Department of Biotechnology and Biophysics, University of Würzburg, 97074 Würzburg, Germany

## Abstract

Traditionally, peripheral sensory neurons hold the monopole of transducing external stimuli. Current research moves epidermal keratinocytes into focus as sensors and transmitters of nociceptive and non-nociceptive sensations, tightly interacting with intraepidermal nerve fibers at the neuro-cutaneous unit. In animal models, epidermal cells establish close contacts and ensheath sensory neurites. However, ultrastructural morphological and mechanistic data examining the human keratinocyte-nociceptor interface are sparse. We investigated this exact interface in human skin applying super-resolution array tomography, expansion microscopy, and structured illumination microscopy. We show keratinocyte ensheathment of nociceptors and connexin 43 plaques at keratinocyte-nociceptor contact sites in healthy native skin. We further derived a fully human co-culture system, modeling ensheathment and connexin 43 plaques *in vitro*. Unraveling human intraepidermal nerve fiber ensheathment and interaction sites marks a milestone in research at the neuro-cutaneous unit. These findings are mind-changers on the way to decipher the mechanisms of cutaneous nociception.

## 1. Introduction

Impairment of the thinly-myelinated A-delta and unmyelinated C-nerve fibers may underlie small nerve fiber pathology observed in patients with peripheral (Birklein, 2005; Lacomis, 2002; Üçeyler *et al*., 2013) and central nervous system diseases (Nolano *et al*., 2008; Weis *et al*., 2011). Cutaneous nerve fiber degeneration and sensitization are hallmarks of small fiber pathology, however, the underlying pathomechanisms are unclear (Üçeyler, 2016). The impact of skin cells on nociceptive and non-nociceptive stimulus detection is increasingly recognized (Lumpkin and Caterina, 2007; Stucky and Mikesell, 2021).

Physiologically, keratinocytes are the predominant cell type in the epidermis and actively participate in sensory signal transduction and nociception at the neuro-cutaneous unit (NCU). In animal *in vitro* cell culture models, selective thermal, chemical, or mechanical keratinocyte stimulation led to activation of co-cultured peripheral neurons (Klusch *et al*., 2013; Mandadi *et al*., 2009; Sondersorg *et al*., 2014). Using animal models, nociceptive behavior was induced in mice selectively expressing transient receptor potential vanilloid 1 (TRPV1) in keratinocytes after capsaicin treatment (Pang *et al*., 2015). Mice expressing channelrhodopsin-2 in keratinocytes also displayed pain behavior and intraepidermal nerve fiber (IENF) derived evoked nerve fiber action potentials during laser stimulation (Baumbauer *et al*., 2015).

For underlying functional stimulus transduction of keratinocytes and IENF, signaling molecules such as adenosine triphosphate (ATP) are increasingly recognized (Mandadi *et al*., 2009; Moehring *et al*., 2018). Hemichannels or gap junctions formed by connexins and pannexins, or vesicular transport may conduct ATP signaling towards afferent nerve fibers (Barr *et al*., 2013; Maruyama *et al*., 2018; Sondersorg *et al*., 2014). Signaling might happen via specialized synapse-like connections to IENF (Talagas *et al*., 2020b). In *Danio rerio* and *Drosophila* models, nerve endings are frequently ensheathed by epidermal cells (Jiang *et al*., 2019; O’Brien *et al*., 2012) and tunneling of fibers through keratinocytes in human skin has recently been shown via confocal microscopy (Talagas *et al*., 2020a). However, the exact mechanisms and mode of signal transduction at the NCU remain elusive.

In an embryonal stem cell-derived 2D human cell culture model, physical contacts between sensory neurons and keratinocytes were observed hinting towards close coupling (Krishnan-Kutty *et al*., 2017). Still, direct and systematic information on ensheathment of human IENF is scarce and ultrastructural architecture or molecular processes remain obscure. Deciphering these contact zones in the human system may profoundly change the understanding of somatosensory processing in health and disease. Ultimately, altered keratinocyte signal molecule release or dysfunctional signaling sites may contribute to cutaneous pain perception (Talagas *et al*., 2017), which could open novel avenues for treatment of small fiber pathology and neuropathic pain (Keppel Hesselink *et al*., 2017).

We aimed at studying exactly these contact zones between keratinocytes and IENF at the NCU in the human system to gain insights on ultrastructure and potential crosstalk in the epidermis. A correlative light and electron microscopy approach via super-resolution array tomography (srAT) in high-pressure frozen, freeze substituted samples (Markert *et al*., 2017) and expansion microscopy (ExM) (Tillberg *et al*., 2016) in diagnostically relevant paraformaldehyde-fixed tissue sections revealed ensheathment and pore protein connexin 43 (Cx43) plaques in native human skin. We further succeeded to establish a fully human keratinocyte and sensory neuron co-culture, which inherited both features of the NCU. We propose a crucial role of nerve fiber ensheathment and Cx43-based keratinocyte-fiber contacts in neuropathic pain and small fiber pathology widening the scope of somatosensorics to non-neuronal cells.

## Materials and methods

### Participants

Healthy volunteers were recruited at the Department of Neurology, University of Würzburg, Germany. For srAT, a 2-mm skin punch biopsy was taken from the back at th10 level (device by Stiefel GmbH, Offenbach, Germany) under local anesthesia following a standard procedure (Üçeyler *et al*., 2010). Tissue sections for ExM and cell cultures were acquired from 6- mm skin biopsy samples taken from the upper thigh according to a previously published protocol (Karl *et al*., 2019). Our study was approved by the Würzburg Medical School Ethics committee (#135/15).

### srAT sample preparation

Biopsies were immediately wetted in freezing solution composed of 20 % (w/v) polyvinylpyrrolidon in phosphate buffered saline (PBS) (0.1 M, pH = 7.4) to prevent dehydration. The epidermal layer was manually dissected from dermal and subdermal compartments of the skin sample and transferred into a type A aluminium specimen carrier (Leica Microsystems, Wetzlar, Germany) with recesses of 200 µm containing polyvinylpyrrolidon and capped with a second carrier without recess (Leica Microsystems, Wetzlar, Germany). Subsequent high pressure freezing and freeze substitution was applied as described previously (Markert *et al*., 2016). 100-nm serial sections were cut via a histo Jumbo Diamond Knife (DiATOME, Biel, Switzerland) with an ultra-microtome EM UC7 (Leica Microsystems, Wetzlar, Germany). Sections were held together as array by adhesive glue (pattex gel compact, Henkel, Düsseldorf-Holthausen, Germany), mixed with xylene (AppliChem, Darmstadt, Germany) and Spinel Black 47400 pigment (Kremer pigmente, Aichstetten, Germany), which was added to the lower side of the LR-White block prior to cutting. Ribbons were collected on poly-L-lysine coated slides (Thermo Fisher Scientific, Waltham, MA, USA).

### srAT immunolabeling, fluorescence imaging, and image processing

Primary and secondary antibodies used for srAT experiments are listed in Table 1. Ultrathin serial tissue sections were encircled via a pap pen (Science Services, München, Germany). A blocking solution containing 0.05 % (v/v) tween20 and 0.1% (w/v) bovine serum albumin in PBS was added for 5 min. Primary antibodies diluted 1:400 in blocking solution were then dropped onto the slides while the initial solution was withdrawn by applying filter paper on the adjacent side of the encircled area. Primary antibodies were incubated for 1 hour in closed humid chambers at room temperature (RT). Samples were washed four times with PBS in 5 min intervals. Afterwards, secondary antibodies were applied for 30 min at RT at 1:400 dilution in blocking solution containing 1:10,000 4’,6-diamidino-2-phenylindole (DAPI; Sigma Aldrich, St. Louise, MO, USA) in the closed humid chamber. Samples were washed again four times with PBS and a last washing step with double distilled H_2_O (ddH_2_O) for 5 min was added. Slides were dried with filter paper, mounted in mowiol 4-88 (Roth, Karlsruhe, Germany), and covered with high precision cover glass No. 1.5H (Roth, Karlsruhe, Germany).

**Table 1:**
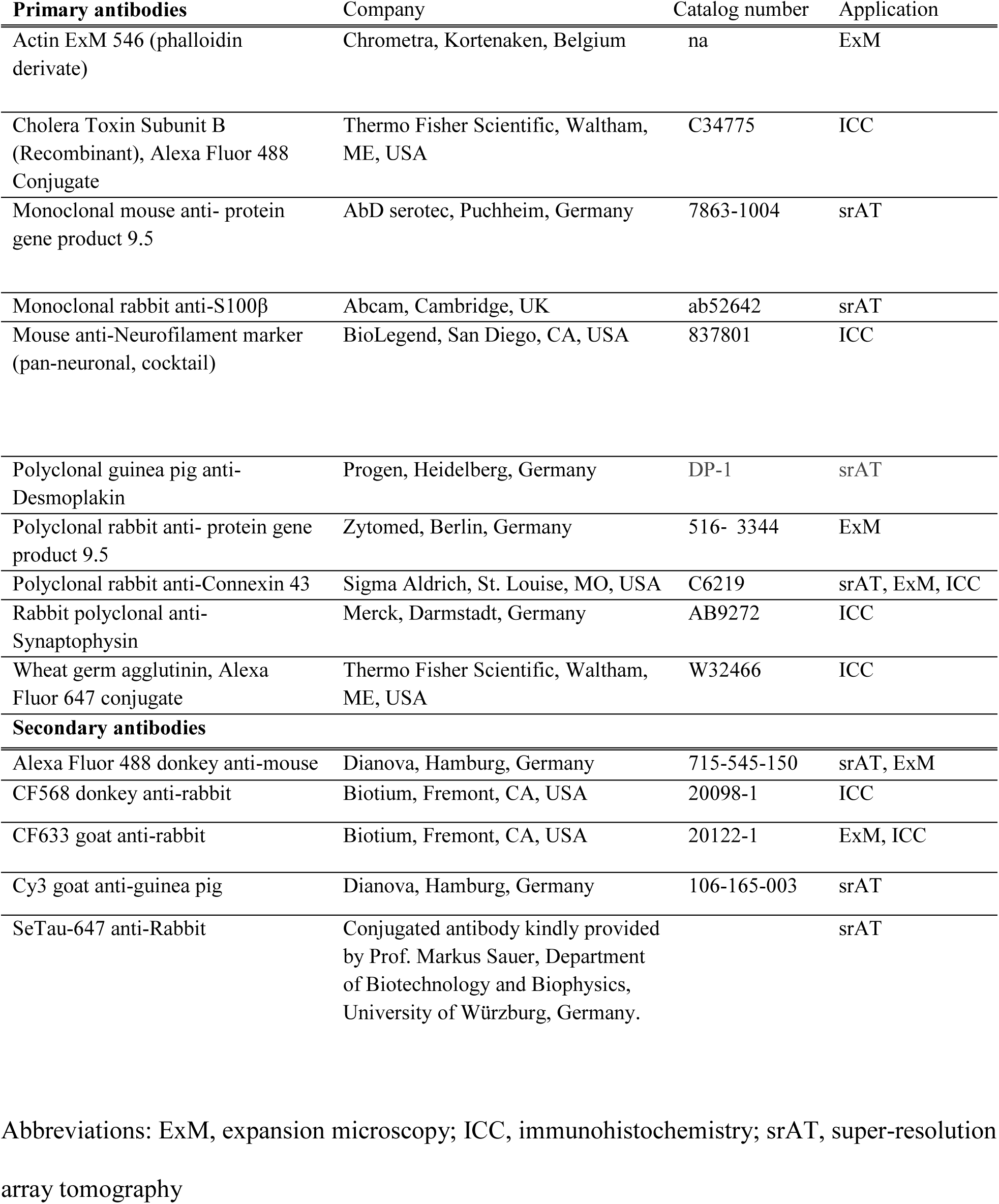
Antibodies and directly conjugated markers. Used reagents in each experiment are indicated under the column ‘Application’.

Image acquisition was performed with the Zeiss ELYRA S.1 SR-SIM with 63x oil-immersion objective plan-apochromat 63x, NA 1.4 Oil Dic M27 and ZEN (black edition) software (all Zeiss, Oberkochen, Germany) with PCO Edge 5.5 sCMOS camera (PCO, Kelheim, Germany), using three rotations. 700 nm z-stacks in 100 nm increments around the observed focal point of DAPI staining per section were imaged. Fluorescence images were processed via image J (version 1.51n, National Institute of Health, Bethesda, MD, USA). Channels were assigned to a defined color code and minimum and maximum of the image histogram adjusted for each channel separately.

The image slice with the brightest signal and best focus within the z-stack was determined for each channel separately to adjust for different light emission wavelength of fluorophores. Each channel was exported as a portable network graphic (png) format file.

### Electron microscopy sample preparation and imaging

For contrasting and carbon coating of the samples, cover glasses were removed and mowiol was washed out with ddH_2_O and blow dried. The object glass area containing the ribbon was cut out with a diamond pen (Roth, Karlsruhe, Germany). A 2.5% (w/v) uranyl acetate (Merck, Darmstadt, Germany) in ethanol solution was dropped onto the sections and incubated for 15 min at RT. Sections were briefly washed in 100% ethanol, 50% (v/v) ethanol in ddH_2_O, and 100% ddH_2_0, followed by 10 min incubation at RT with 50% (v/v) lead citrate solution in decocted H_2_O containing 80 mM lead citrate (Merck, Darmstadt, Germany) and 0.12 M trisodium citrate (AppliChem, Darmstadt, Germany) (Reynolds, 1963). After washing in ddH_2_O, sections were dried and attached to specimen pin mounts via carbon conductive tape (Plano, Wetzlar, Germany). Conductive silver (Plano, Wetzlar, Germany) was applied, connecting the glass with the edges of the holder. A 5-nm carbon coat was applied, using a CCU-010 carbon coating machine (Safematic, Bad Ragaz, Switzerland). Serial sections were imaged in a JSM- 7500F field emission scanning electron microscope (SEM; JEOL, Tokyo, Japan) with an acceleration voltage of 5 kV, a probe current of 0.3 nA, and a working distance of 6.0 mm. At each area of interest, several images with increasing magnification were acquired.

### srAT image processing, correlation, and modelling

Montage and alignment of scanning electron microscopy (SEM) images were achieved via the ImageJ plugin TrakEM2 (version 1.0a, 04.07.2012) (Cardona *et al*., 2012; Schindelin *et al*., 2012). Images corresponding to the same section at different magnifications were merged within one layer with least squares montage in similarity mode and an alignment error of 10-20 pixel. After each z-layer was positioned, serial 100-nm sections were orientated via align layers, using similarity as transformation mode and 20-100 pixel alignment error. The area of interest in each layer was exported in a tagged image file (tif) format. To correlate immunofluorescence (IF) and SEM information, associated IF channel images and montaged SEM images were loaded into the vector graphics editor program Inkscape (version 0.92.3, 11.03.2018) and processed according to a standardized protocol (Markert *et al*., 2017). IF channel images were overlaid and linked, leaving only the DAPI channel visible as first image layer. Opacity of IF images was reduced and DAPI labeled heterochromatin was used as an independent and unbiased landmark for correlation. Linked IF images were linearly transformed (rotation and resizing, but no distortions) to fit the cell nuclei orientation of the EM image. When adequate overlay was achieved, a rectangular area containing the region of interest (ROI) was extracted and each layer exported as a png file. Corresponding IF and EM images were then imported into the image editor GIMP2 (Version 2.10.0, 02.05.2018) for appropriate overlay and exported as png files. For tracing IENF in 3D, the open source software package IMOD was used (Kremer *et al*., 1996). Alternating 100-nm sections were imaged via IF and correlated with their corresponding EM images. Within the 100-nm stepwise srAT stack, the trajectory of an IENF was volumetric reconstructed as extrapolated tubular structure. Its position was determined based on PGP9.5 localization available for every second section. Furthermore, distinguishable electron density compared to keratinocyte cytoplasm and absence of desmosomes between adjacent keratinocytes were considered to identify the IENF in the EM context.

### Skin cryosections

PFA fixed 10-µm skin cryosections were blocked in 10% BSA(w/v) in PBS for 30 min and incubated with primary antibodies against 1:100 PGP9.5 and Cx43 in 0.1% (w/v) saponin and 1% (w/v) BSA in PBS over night at 4°C.Applied antibodies are listed in Table 1. After washing with PBS, secondary antibodies were applied for 2 h at RT with 1% BSA(w/v) in PBS. After washing, sections were covered with droplets of PBS and stored at 4°C until further processing.

### Expansion microscopy

ExM was adapted from former published protocols (Tillberg *et al*., 2016; Zhao *et al*., 2017). Skin sections were incubated with PBS containing 0.1 mg/ml Acryloyl-X (Thermo Fisher Scientific, Waltham, MA, USA) in dimethyl-sulfoxide (Sigma Aldrich, St. Louis, MO, USA) over night at RT. Afterwards, 33 nM expandable phalloidin derivate Actin ExM 546 (Chrometra, Kortenaken, Belgium), labeling actin cytoskeleton, was applied in PBS with 1% (w/v) BSA and 0.1 mg/ml Acryloyl-X for 1 h at RT. Subsequently, a monomer solution containing 8.625% (w/w) sodium acrylate (Sigma-Aldrich, St. Louis, MO, USA), 2.5% (w/w) acrylamide (Sigma-Aldrich, St. Louis, MO, USA), 0.15% (w/w) N,N‘-methylenbisacrylamide (Sigma-Aldrich, St. Louis, MO, USA), and 11.7% (w/w) sodium chloride (Sigma-Aldrich, St. Louis, MO, USA) in PBS was added at 4°C for 30 min. Gelation was performed after replacement with fresh monomer solution, additionally containing 0.2% (w/v) ammonium persulfate (Sigma-Aldrich, St. Louis, MO, USA), 0.2% (v/v) tetramethylethylenediamine (Sigma-Aldrich, St. Louis, MO, USA), and 0.01% 4- Hydroxy-TEMPO (w/v) (Sigma-Aldrich, St. Louis, MO, USA). Sections were first incubated at 4°C for 30 min followed by 2h at 37°C. A gelation chamber assembled with each two coverslip pieces No1 (R. Langenbrinck, Emmendingen, Germany) on the side as spacers and one on top, serving as a lid, in a humidified plastic chamber was used to enable uniform gelation. Gelated samples were digested in 4 U/ml proteinase K buffer (New England Biolabs, Ipswitch, MA, USA) with 50 mM Tris pH 8.0 (Serva, Heidelberg, Germany), 50 mM EDTA (Sigma-Aldrich, St. Louis, MO, USA), 0.5% (v/v) Triton X-100 (Thermo Fisher Scientific Scientific, MA, USA) and 0.8 M guanidine HCl (Sigma-Aldrich, St. Louis, MO, USA) for 2 h at 60°C. Subsequently, gels were washed 10 min at RT with PBS, then with 1:2,500 DAPI (Sigma Aldrich, St. Louise, MO, USA) in PBS for 20 min at RT and again 10 min in BPS at RT. Gels were transfer into a dark petri dish with 100x times final gel volume of sterile ddH_2_O with a razor blade. Gels were expanded for at least 1 h at RT before direct post-expansion imaging or storage at 4°C.

Labeled sections were imaged both in pre-expansion and post-expansion state with an DMi8 inverse microscope via 20x dry objective HC PL FLUOTAR L 20x/0.40, 11506243, LAS X software, and DMC300G monochrome camera (all Leica Microsystems, Wetzlar, Germany) to determine the expansion factor via manual alignment in inkscape. Further, imaging was performed using the ELYRA S.1 SR-SIM with 63x water-immersion objective C-Apochromat, 63 x 1.2 NA, 441777-9970 (all Zeiss, Oberkochen, Germany), and a PCO Edge 5.5 sCMOS camera (PCO, Kelheim, Germany). Gels were imaged inside poly-D lysine (Sigma Aldrich, St. Louise, MO, USA) coated imaging chambers (Thermo Fisher Scientific, Waltham, ME, USA) to prevent drifting. Non-computed (widefield) images were used. Min/Max values were processed via ImageJ (version 1.51n, National Institute of Health, Bethesda, MD, USA) for visualization.

### Fully human co-culture system

Human induced pluripotent stem cells (iPSC) derived from fibroblasts were differentiated into sensory neurons of a healthy control cell line as previously described (Klein et al., submitted). Co-culture chambers (ibidi, Gräfelfing, Germany) were attached to 12-mm BioCoat® Poly-D-Lysine/Laminin coverslips (Corning, New York, NY, USA). Both inner chambers were additionally coated with 1:50 matrigel growth factor reduced (Corning, Corning, New York, NY, USA) at 37°C for 30 min. Four-week old neurons were detached via TrypLE (Thermo Fisher Scientific, Waltham, MA, USA), transferred into falcon tubes containing DMEM/F12 (Dulbecco’s Modified Eagles Medium/Nutrient Mixture F-12; Thermo Fisher Scientific, Waltham, MA, USA) at 37 °C, and centrifuged for 3 min with 500 x g at RT.

Conditioned neuronal medium, consisting of DMEM/F12 GlutaMAX + 1X B-27 Plus Supplement + 1X N-2 Supplement + 100 U/ml 1% penicillin/ streptomycin (pen/strep; all Thermo Fisher Scientific, Waltham, MA, USA) + 20 ng/ml BDNF + 20 ng/ml GDNF + 20 ng/ml NGFb (all Peprotech, Rocky Hill, NJ, USA) + 200 ng/ml ascorbic acid (Sigma-Aldrich, St. Louis, MO, USA), spiked with 10 µM floxuridine (Santa Cruz Biotechnology, Dallas, TX, USA), was withdrawn prior to TrypLE treatment, filtrated via 0.2 µm syringe filters (Sarstedt, Nümbrecht, Germany) and kept at 37°C. Neurons were resuspended in 70 µl of filtered conditioned neuronal medium and seeded into one chamber compartment and acclimated for one week. Healthy control-derived primary keratinocytes were acquired and cultured via routine methods (Karl *et al*., 2019) and seeded into the corresponding compartment. The chamber insert barrier separating the associated chambers was removed after 24 h and medium exchanged either with fresh keratinocyte medium, comprising of EpiLife Medium supplemented with 1% EpiLife defined growth supplement, and 1% pen/strep (all Thermo Fisher Scientific, Waltham, USA) or stored conditioned neuronal medium. After 6 days, co-cultures were fixed with 4% PFA (v/v) in PBS ^Ca++ / Mg++^ (PBS^++^) at RT for 15 min and washed three times 5 min in PBS. Briefly, coverslips were treated with10 μg/ml wheat germ agglutinin, conjugated with Alexa Fluor 647 (WGA-647; Thermo Fisher Scientific, Waltham, MA, USA) in PBS for 10 min at RT and washed two times with PBS. Subsequently cells were blocked 30 min with 10% FCS (v/v) and 0.1% (w/v) saponin in PBS^++^, then labeled either 1:100 anti-Cx43, or 1:100 anti-synaptophysin antibodies for visualization of contact sites over night at 4°C. Antibody solution contained 10% FCS (v/v) and 0.1% saponin (w/v) in PBS. After washing, secondary antibodies, 1:10,000 DAPI, and 10 μg/ml cholera toxin subunit B, conjugated with Alexa Fluor 488 (Thermo Fisher Scientific Scientific, MA, USA) were applied for 30 min at RT in antibody solution without saponin. Coverslips were transferred onto object holders, embedded with mowiol 4-88 (Roth, Karlsruhe, Germany) and -stored at 4°C until further processing.

### 2D co-culture live imaging and fluorescence microscopy

Directly before barrier removal, the co-cultures were transferred into a Lab-Tek chamber system (Thermo Fisher Scientific, Waltham, MA, USA) with two-well compartments. The remaining well was filled with 500 µl PBS^++^. After barrier removal and addition of conditioned neuronal medium, co-cultures were incubated for 2d. For live-imaging, the Lab-Tek slide was transferred to an inverse DMi8 microscope operated on LAS X software (Leica Microsystems, Wetzlar, Germany), equipped with a live imaging chamber (ibidi, Gräfelfing, Germany). Phase contrast images with 20x objective HC PL FLUOTAR L dry 20x/0.40, 11506243 were taken in 20 min intervals for 67 h with a DMC2900 color camera (both Leica Microsystems, Wetzlar, Germany), with cells kept at 37°C, with 5% CO_2_, and 20% O_2_ (both v/v). Single regions were stitched together as overview via Fiji plugin MosaicJ (Thévenaz and Unser, 2007). Min/Max values were processed via ImageJ (version 1.51n, National Institute of Health, Bethesda, MD, USA) for visualization.

Fluorescently labeled co-cultures were imaged with an Axio Imager.M2 (Zeiss, Oberkochen, Germany), equipped with spinning disc-confocal system (X-light V1, CrestOptics, Rome Italy) and Spot Xplorer CCD camera (SPOT Imaging, Sterling Heights, MI, USA) operated on VisiView software (Visitron Systems, Puchheim, Germany) for overview and Lattice-SIM for detailed super-resolution analysis with 63x water immersion C-Apochromat 63x/1.2 W Korr UV-VIS-IR M27, 21787-9971-790 and ZEN (black edition) software (all Zeiss, Oberkochen, Germany), with two aligned PCO Edge 4.2 M sCMOS cameras (PCO, Kelheim, Germany). Min/Max values were processed via ImageJ (version 1.51n, National Institute of Health, Bethesda, MD, USA) for visualization. For 3D visualization, Min/Max values of complete z-stacks were adjusted per channel in ZEN (blue edition) software (Zeiss, Oberkochen, Germany) and depicted in 3D mode.

## Results

### Human intraepidermal nerve fiber segments are engulfed by keratinocytes

While many IENF were passing between neighboring keratinocytes, srAT revealed IENF ensheathed by keratinocytes in all three healthy subjects. This tunneling of fibers was observed both in the basal and upper epidermal layers for several consecutive 100-nm thin sections indicated via PGP9.5 labeling (Figure 1A). IENF intersected closely to either the lateral, posterior, or anterior boundary of the respective keratinocyte and was engulfed by the respective cell for several µm (Figure 1A, Video1). For advanced tracing of an IENF through the epidermis, a representative site was reconstructed (Figure 1B, Video2). Whilst a major part of the respective nerve fiber grew in close contact to and in between keratinocytes, a substantial portion tunneled through one basal keratinocyte. Specificity of antibodies was examined by tracking the fluorophore signal within consecutive slices and negative control via omission of the primary antibody (Figure supplement 1). High pressure freezing and freeze substitution followed by LR- White embedding preserved the ultrastructure and antigenicity of the human skin tissue well, illustrated by identification of collagen fibrils, desmosomes, nuclei, and mitochondria (Figure 1C).

**Figure 1.**
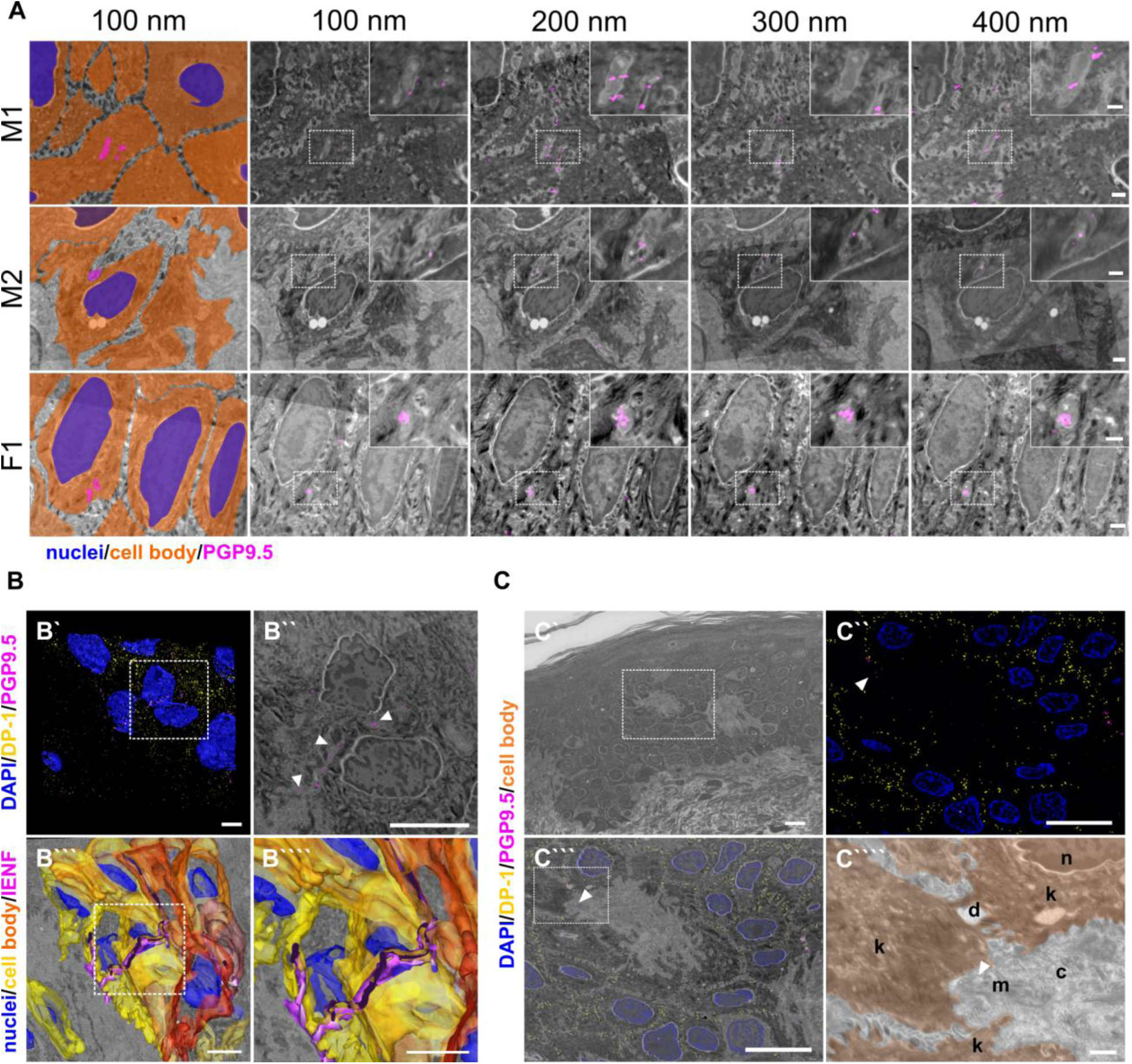
Epidermal nerve fiber ensheathment and srAT functionality. (A) Ensheathment of IENF by keratinocytes. Nerve fibers projecting within keratinocytes in skin punch biopsy samples of two male (M1, M2) and one female subject (F1). First tile shows keratinocyte cell bodies (orange), nuclei (blue), and fiber (magenta) in pseudo color. Each row represents four consecutive sections with 100 nm thickness of correlated images, with PGP9.5 labeling for IENF (magenta), while dashed insets show higher magnification of the region of interest in inlay. See also Video1. Scale bar 1 µm, magnified insets 500 nm. (B) 3D reconstruction of IENF processes traversing between and within keratinocytes. (B’) 3D visualization of fluorescence signal from srAT approach, white rectangle indicates area in b’’. PGP9.5 (magenta) labeled nerve fiber processes between and in keratinocytes in close apposition to nuclei (blue). DP-1 (yellow) marks intercellular desmosomal junctions as keratinocyte cell boundaries. (B’’) Single plane with overlay of PGP9.5 signal and EM. (B’’’) Extrapolation of IENF trajectory in 3D, based on IF signal and EM ultrastructure with fiber (magenta), keratinocyte cell bodies (yellow-orange), and nuclei (blue); see also Video2. Scale bars 5 µm. (C) Preservation of antigenicity and cellular structure in LR-White embedded epidermal tissue with Overview area from SEM (C’). Scale bar 10 µm. (C’’) SIM image of 100-nm skin section with DP-1 (yellow), PGP9.5 (magenta), and DAPI (blue) labeling. Arrowhead indicates PGP9.5- positive IENF processes. Scale bar 10 µm. (C’’’) Correlated SIM and SEM image from dashed rectangle in A. Scale bar 10 µm. (C’’’’) Inset of c showing subcellular preservation of collagen fibers (co), desmosomes (de), keratinocytes (k), mitochondria (m), and nucleus (n). Arrowhead indicates IENF processes also observed via IF in a. Scale bar 1 µm. Abbreviations: DP-1, desmoplakin 1; IENF, intraepidermal nerve fiber; IF, immunofluorescence; LR-White, London Resin-White; PGP9.5, protein gene product-9.5; SEM, scanning electron microscopy; SIM, structured illumination microscopy.

### srAT facilitates tracing of further skin cell populations

The combination of cellular markers such as PGP9.5 or S100β with information on cell morphology from EM scans allowed tracing of various cell populations via srAT in human skin. S100β-positive Langerhans cells were visualized in the epidermis and their dendritic protrusions were traced between keratinocytes (Figure 2A and B). Within the dermis, S100β-PGP9.5-co- localized signal identified dermal Schwann cells enwrapping nerve fiber processes (Figure 2C-D).

**Figure 2.**
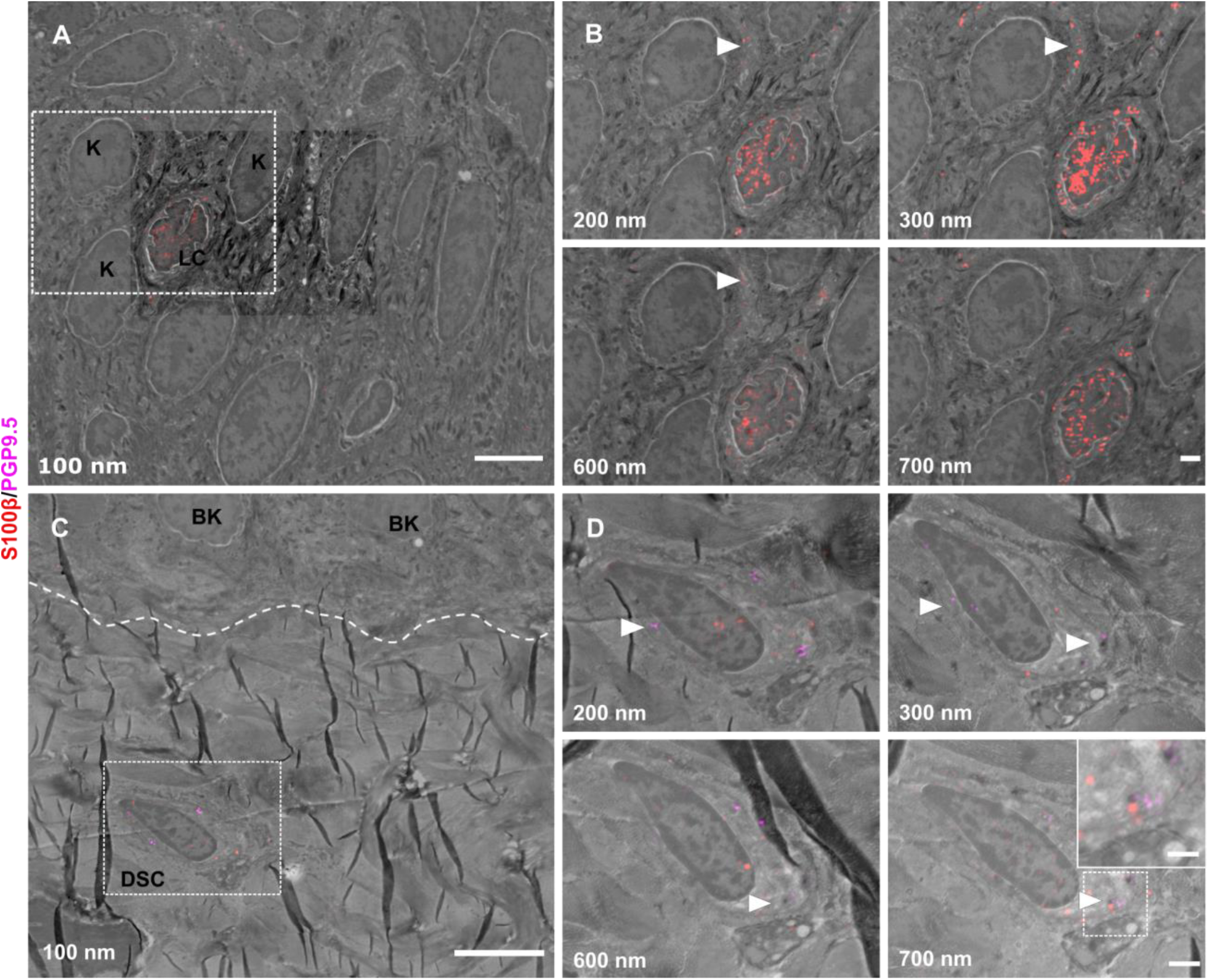
General utility of srAT for tracing skin cells. (A) Epidermis with keratinocytes (K) and Langerhans cell (LC) (B) Single Z-sections of outlined areas. LC was identified via S100β (red). Arrowheads indicate LC protrusions in contact to K. (C) Upper dermis and basement membrane (dashed line) with basal keratinocytes (BK) and dermal Schwann cell (DSC). (D) Single Z-sections of outlined areas. DSC was identified via PGP9.5 (magenta) and S100β (red) labeling. Dermal fiber processes are enwrapped by CSC, indicated with arrowheads. Inset in last panel shows magnification of marked fiber. Scale bars 5 µm (A, C), 1 µm (B, D), and 500 nm (inset in D) Abbreviations: BK, basal keratinocyte; DSC, dermal Schwann cell; K, keratinocyte; LC, Langerhans cell; PGP9.5, protein product 9.5; S100β, S100 calcium binding protein B.

### Nerve fiber ensheathment can be visualized by ExM in diagnostic skin samples

To visualize ensheathed nerve fibers also in thicker, diagnostically used skin punch biopsy sections, we applied ExM allowing super-resolution imaging with epifluorescence microscope setups. Expansion factor of samples fell between 4.3-4.4x and showed isotropic epidermal expansion as documented via pre and post-expansion acquired images (Figure 3). Actin filaments were used as a marker to outline epidermal cell bodies and IENF entry and exit points, while cytoplasmic PGP9.5 identified IENF. Nerve fibers were traced via ExM for their course through the epidermis, being ensheathed over several µm (Video3).

**Figure 3.**
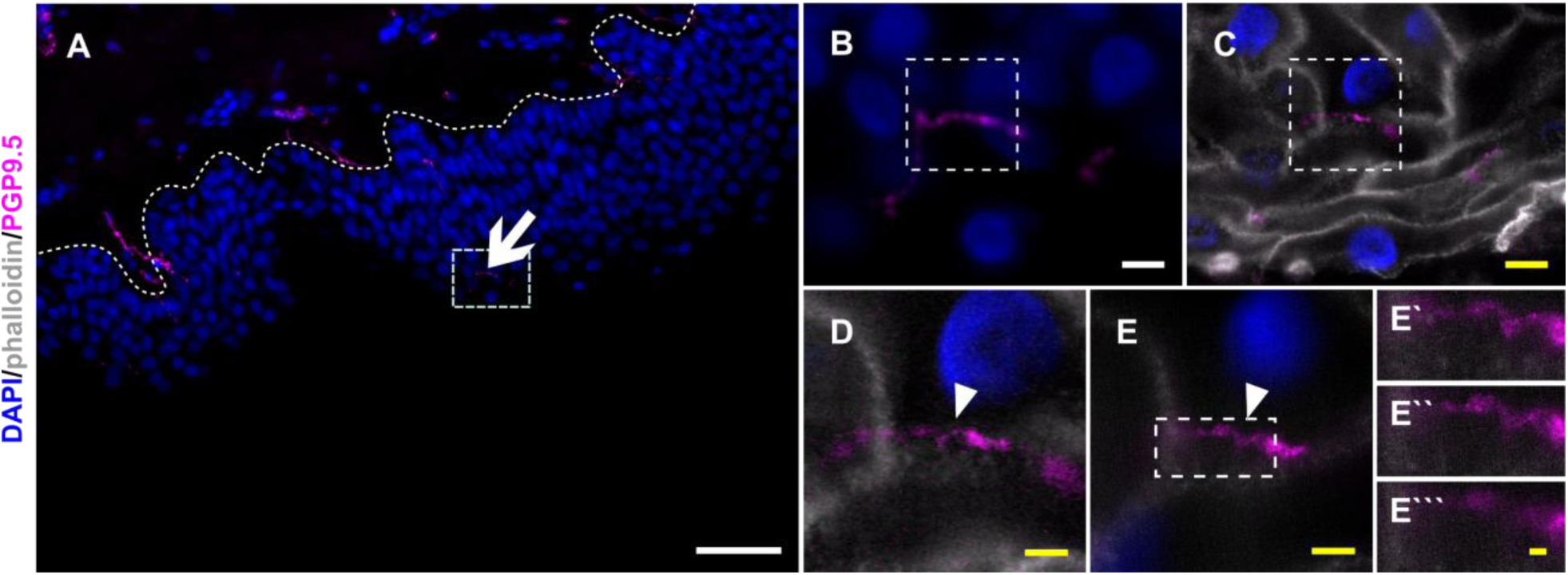
Resolving nerve fiber ensheathment in expanded skin tissue. (A) Overview of skin section prior to expansion with PGP9.5-positive IENF (magenta) and nuclear DAPI (blue) signal; dotted line illustrated epidermis-dermis border and arrow indicates IENF. White rectangle marks inset enlarged in B. (B) Enlarged area prior to expansion and (C) matched area post-expansion at same magnification with addition of actin marker phalloidin (grey), white rectangles mark inset area of D and E.(D) enlarged IENF area at 20x magnification and (E) at 63x magnification with arrowheads indicating IENF passing through keratinocyte. Inset in E marks enlarged area in È-È’’. (È) shows z-plane prior to E, (È’) same z-plane as E, and È’’ z-plane step after E. White scale bars indicate pre-expansion state, yellow scale bars were corrected for expansion factor. Scale bars: 50 µm (A), 5 µm (B, C), 2µm (d, e), 500 nm (È’’). Z-step size of 1.2 µm translates to approximate 276 nm in expanded gel. Abbreviations: Cx43, connexin 43; DAPI, 4’,6-diamidino-2-phenylindole; IENF, intraepidermal nerve fiber; PGP9.5, protein gene product-9.5

### Cx43 plaques as potential keratinocyte-nerve fiber communication sites

Connexin hexamers can form hemichannels acting as small pores. In open state, small molecules can pass and be released from the cell, which was already shown for ATP and Cx43 (Weber *et al*., 2004). This may allow purinergic signaling towards neighboring cells or nerve terminals in close proximity. For srAT, Cx43 labeling was assumed a *bona fide* signal, if ≥2 consecutive sections showed fluorescent staining, translating to 200-400 nm. These clusters were mostly found at keratinocyte-keratinocyte contact zones (Figure 4A), however, distinct Cx43 plaques were also identified in direct proximity to single IENF when growing between keratinocytes (Figure 4B).

**Figure 4.**
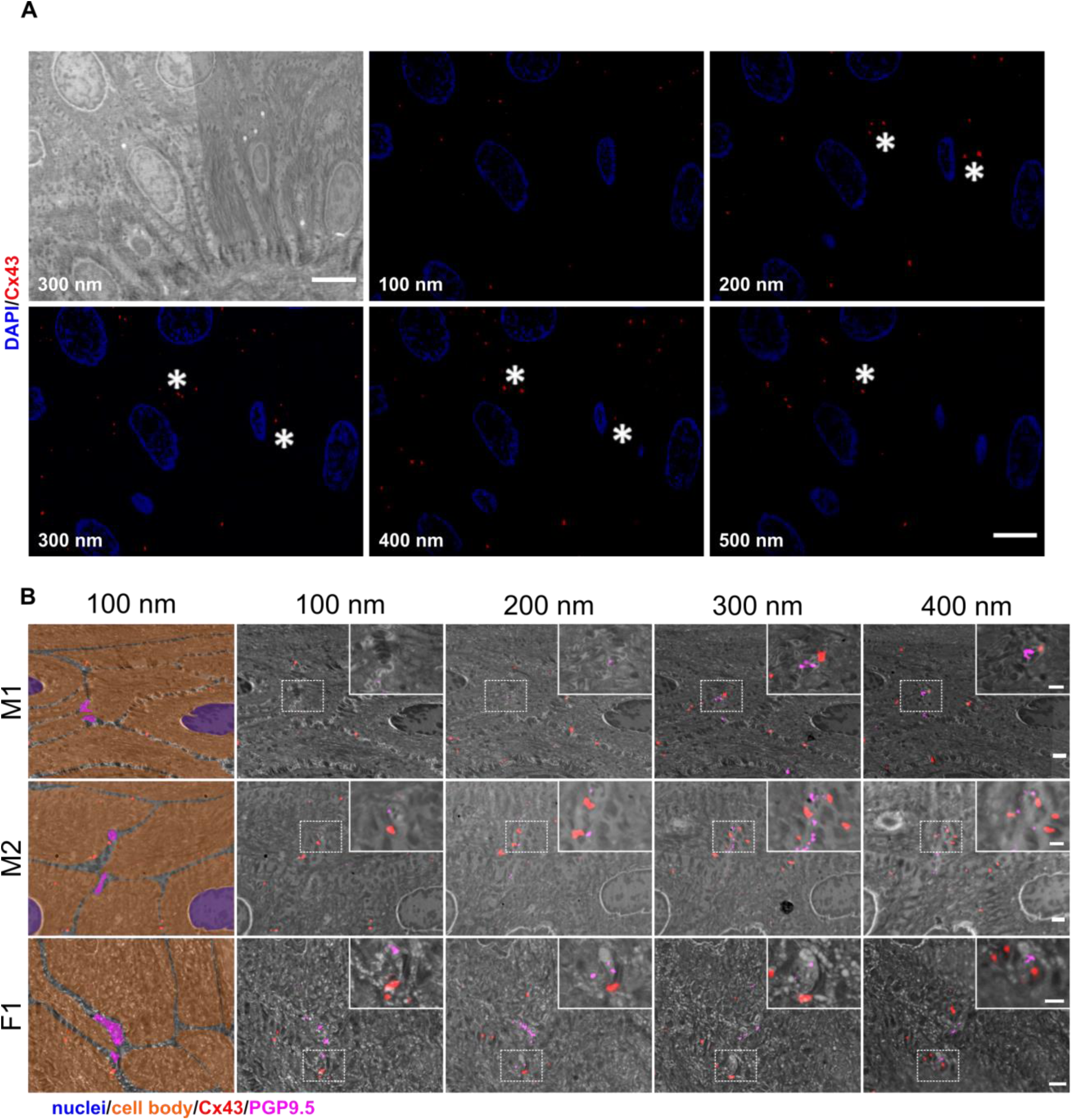
Identification of Cx43 plaques via srAT (A) Tracking of Cx43 plaques in epidermal layers. First panel illustrates SEM overview of epidermal layers corresponding to five consecutive sections of IF images showing Cx43 signal (red) and nuclei (blue). Asterisks show examples of traced Cx43 plaques. (B) Cx43 plaques at keratinocyte-nerve fiber close contact sites. Nerve fibers processing between keratinocytes in skin samples of two male subjects (M1, M2) and one female subject (F1). Each row represents four consecutive sections of 100 nm thickness. First tile shows keratinocyte cell bodies (orange), nuclei (blue) and fiber (magenta), in pseudo color with Cx43 signal (red). Correlated PGP9.5 labeling (magenta) locates at nerve fibers and Cx43 labeling (red) indicates Cx43 plaques. Insets show magnification of contact area. Scale bars: 5 µm (A), 1 µm (B), magnified insets 500 nm. Abbreviations: Cx43, connexin 43; IF, Immunofluorescence; PGP9.5, protein gene product-9.5; SEM, scanning electron microscopy.

In analogy to nerve fiber ensheathment, we investigated Cx43 accumulations also in expanded diagnostic skin samples. In pre-expansion state, the attribution of single Cx43 plaques to specific sites between keratinocytes or towards IENF was hardly possible, due to the compact structure of the epidermis. However, after expansion, specific Cx43-positive accumulations in direct contact to PGP9.5-positive IENF could be identified (Figure 5; Video3).

**Figure 5.**
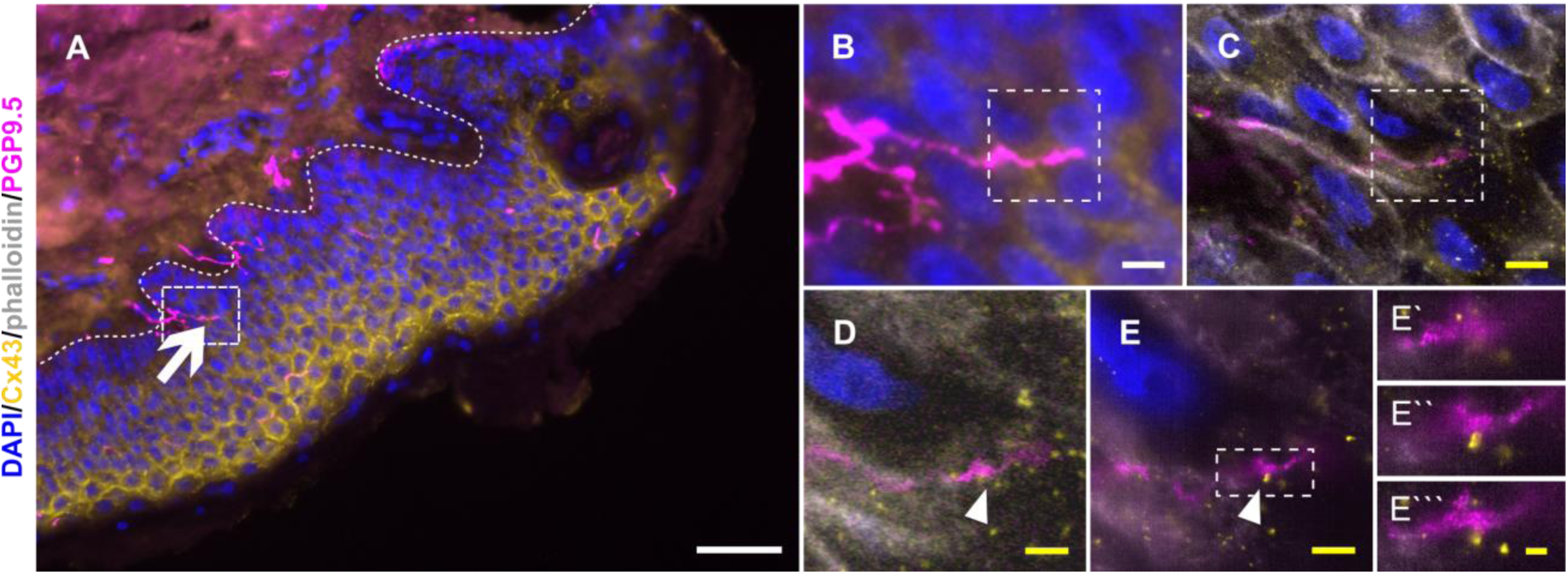
Cx43 accumulation at keratinocyte-nerve fiber contact site in expanded epidermis. (A) Overview of skin section prior to expansion with PGP9.5-labeled IENF (magenta), Cx43 (yellow), and nuclear DAPI (blue); dotted line illustrates epidermis-dermis border and arrow indicates IENF. White rectangle marks inset enlarged in B. (B) Enlarged area prior to expansion and (C) matched area post-expansion at same magnification with addition of actin marker phalloidin (grey), white rectangles mark inset area of D and E.(D) enlarged IENF area at 20x magnification and (E) at 63x magnification with arrowheads indicating Cx43 plaque at IENF. Inset in e marks enlarged area in È-È’’. (È) shows z-plane prior to E, (È’) same z-plane as E, and È’’ z-plane step after E. White scale bars indicate pre-expansion state, yellow scale bars are corrected for expansion factor. Scale bars: 50 µm (A), 5 µm (B, C), 2µm (D, E), 500 nm (È- È’’). Z-step size of 1.2 µm translates to approximate 274 nm in expanded gel. Abbreviations: Cx43, connexin 43; IENF, intraepidermal nerve fiber; PGP9.5, protein gene product-9.5.

### Neurites establish contacts to keratinocytes in fully human co-culture system

Sensory neurons and keratinocytes each formed clusters after seeding into two-comparted chambers (Figure 6A-C). After barrier removal, neurites actively grew towards keratinocytes and established contacts within few days (Figure 6C and D, Video4). Neurite-keratinocyte contacts were apparent, both in conditioned neuronal medium and keratinocyte medium. However, keratinocytes underwent terminal differentiation in neuronal medium, while predominantly maintaining a basal state in keratinocyte medium (Figure supplement 2).

**Figure 6.**
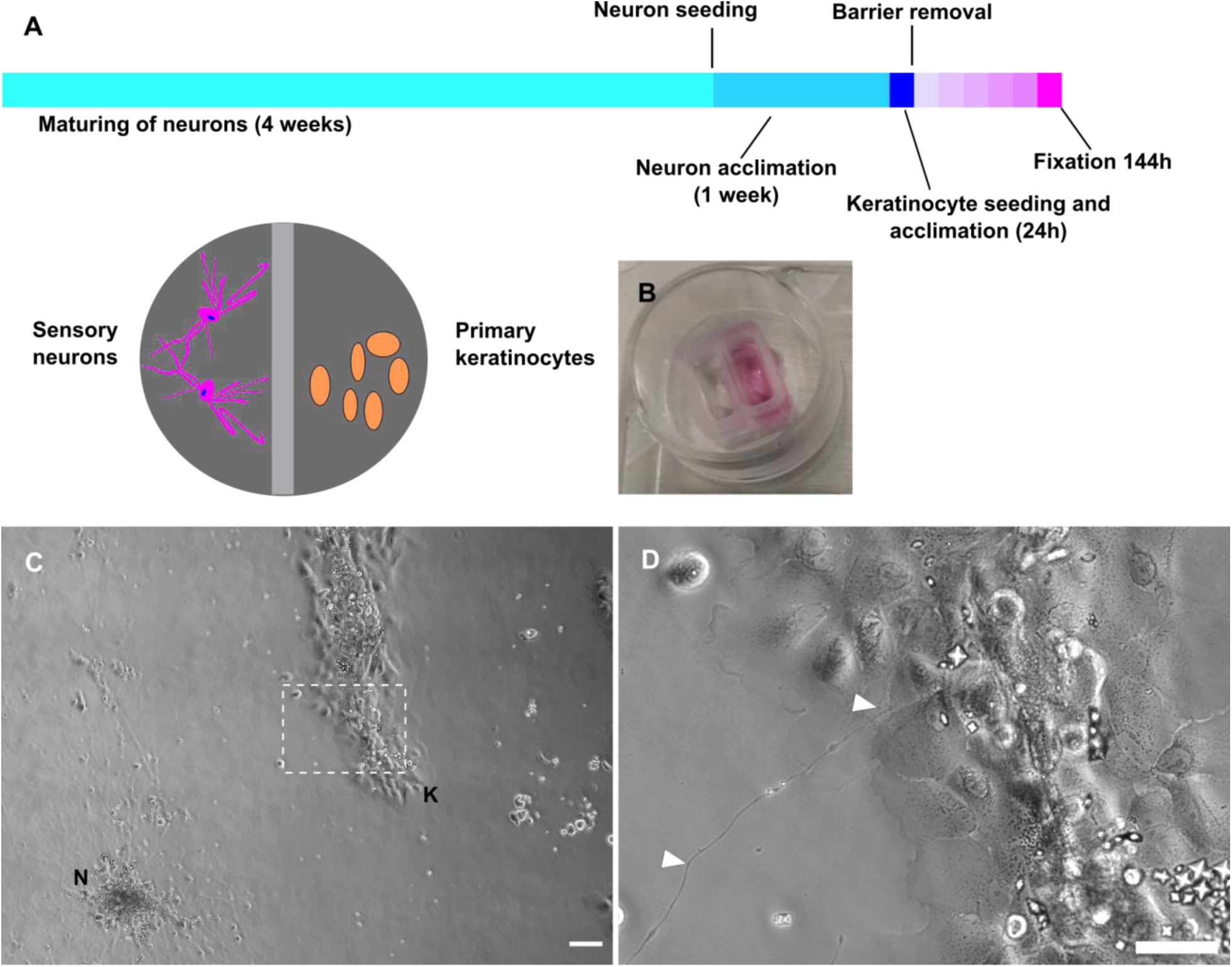
Fully human sensory neuron-keratinocyte co-culture model. (A) Timeline of culturing protocol and compartment scheme. (B) Chamber system. (C) Overview of co-culture after 115h in conditioned neuronal medium with neuronal cluster (N) and keratinocyte colony (K). Inset (D) shows a single neurite in contact with keratinocytes (arrowheads). Co-culture kept in conditioned neuronal medium. Scale bars 100 µm (C), magnified inset 50 µm (D). See also Video4. Abbreviations: K, keratinocyte colony; N, neuronal cluster.

### Ensheathment and Cx43 complexes are present in a fully human co-culture model

In order to distinguish the neurite versus keratinocyte membrane, we specifically labeled sensory neurons via cholera toxin subunit B (Ctx) targeting the ganglioside monosialotetrahexosylganglioside 1 (GM1) (Dederen *et al*., 1994; Tong *et al*., 1999). Conversely, the membrane of keratinocytes was targeted by wheat germ agglutinin (WGA) (Belleudi *et al*., 2011; Watt, 1983). We identified neurite-keratinocyte contacts using confocal microscopy (Figure 7A and B) and observed ensheathment via lattice-SIM super-resolution microscopy (Figure 7 C-C’’). Non-ensheathed neurites frequently passed in close proximity and over keratinocytes (Figure 7D). Intriguingly, Cx43 labeling revealed Cx43 plaques at those passing sites, distributed over several individual keratinocytes (Figure 7D-F’’’).

**Figure 7.**
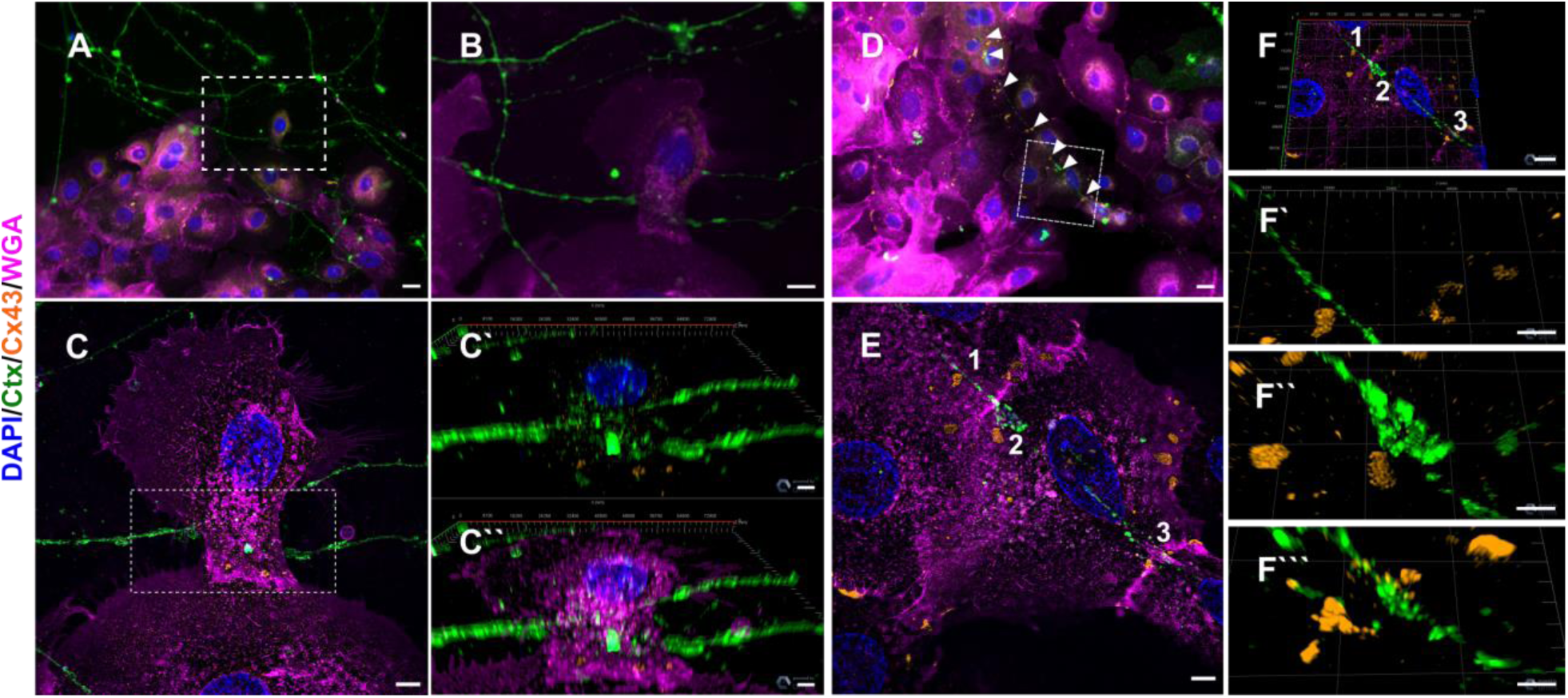
Neurite ensheathment and Cx43 plaques in full human co-culture model. (A) Confocal overview image of Ctx-positive sensory neurites (green), Cx43 positive (orange) keratinocytes with membrane labeling of keratinocytes via WGA (magenta) and nuclear DAPI (blue). (B) Inset of ensheathment area from A. (C) Single plane lattice SIM image and respective inset area with 3D visualization of z-stack (2.925 µm depth, 0.196 µm steps) showing nucleus, Cx43, and neurite signal (C’) and including WGA (C’’). (D) Confocal overview image of Ctx-positive sensory neurites (green), Cx43-positive (orange) keratinocytes with membrane labeling of keratinocytes via WGA (magenta) and nuclei (blue). Arrowheads indicate Cx43 - neurite contact areas. (E) Single plane lattice SIM image and respective inset area with 3D visualization, numbers represent single Cx43 plaques (F) of z-stack (2.925 µm depth, 0.196 µm steps). (F’-F’’’) detailed neurite - Cx43 contact areas. Co-culture kept in keratinocyte medium. Scale bars: 20 µm (A, D), 5 µm (C-C’’, E, F’-F’’’’), 10 µm (F). Abbreviations: Ctx, cholera toxin subunit B; Cx43, connexin 43; DAPI, 4’,6-diamidino-2-phenylindole; SIM, structured illumination microscopy; WGA, wheat germ agglutinin.

We further found neurites growing in a gutter-like structure (Figure 8A), merging into the keratinocyte membrane (Figure 8B) and observed neurites that establish bouton-like contacts with keratinocytes (Figure 8C). We included SYP labeling as a marker for small synaptic vesicles, which might serve as another pathway of signal transduction between keratinocytes and IENF (Talagas *et al*., 2020b). In our iPSC-derived neurons, SYP was distributed throughout the cytoplasm and not restricted to the cytoskeleton (Figure supplement 3). Conversely, only weak SYP labeling, not associated with neurite contact sites, was present in keratinocytes (Figure 8A- C).

**Figure 8.**
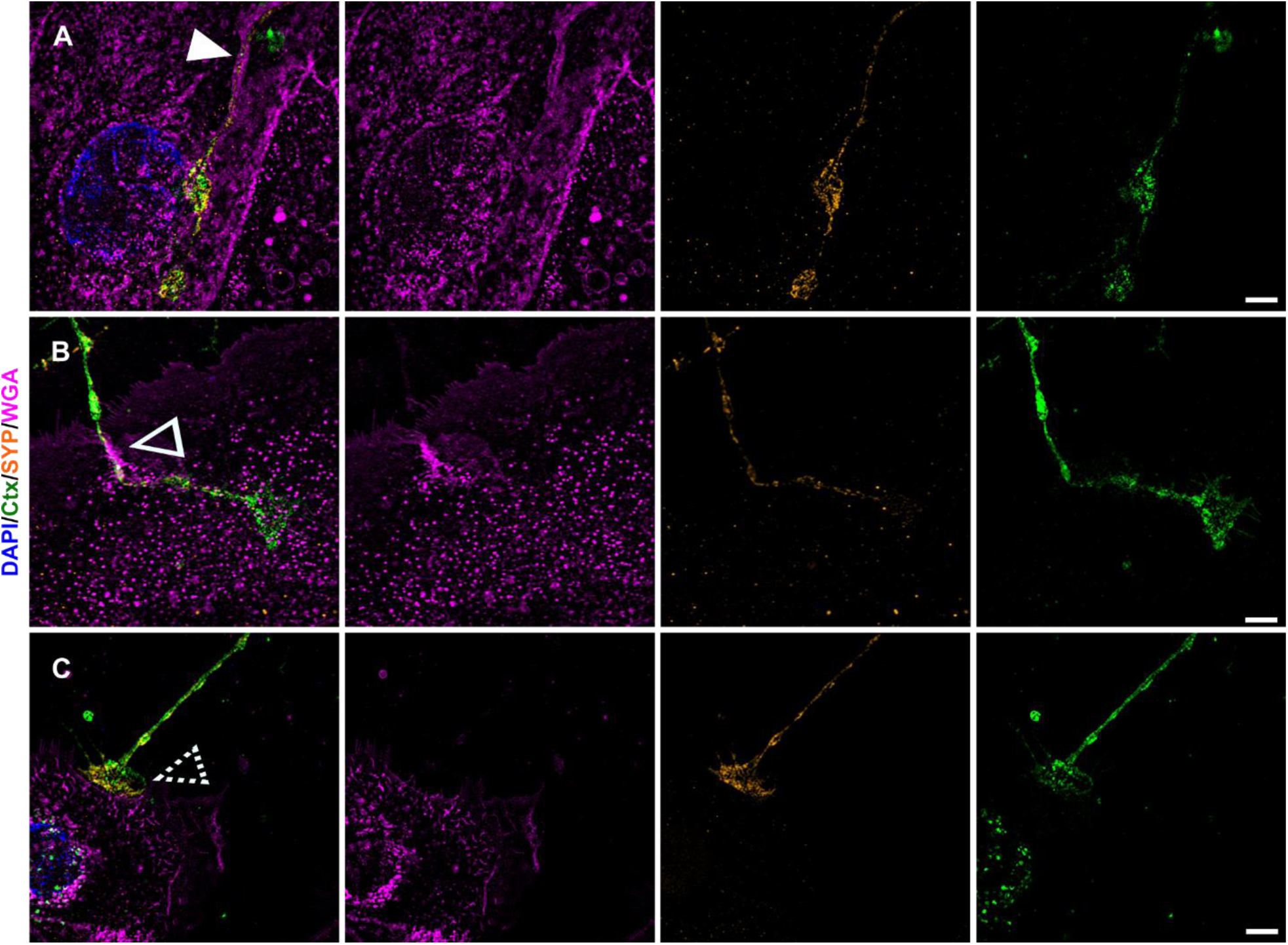
Further keratinocyte-neurite interactions and synaptic vesicular SYP distribution. Single plane lattice SIM images with overlay of nuclear DAPI (blue), WGA (magenta), SYP (yellow), and Ctx (green) signal as first panel, followed by single channel images of WGA, SYP, and Ctx. Distinct contact sites with gutter like structure (A) indicated by filled arrowhead, enwrapping (B) indicated via hollow arrowhead, and bouton-like contact (C) indicated via dashed arrowhead were observed in human co-culture. SYP signal in a-c is predominantly restricted to neurite with sparse dotted labeling in keratinocytes. Co-culture kept in keratinocyte medium. Scale bars: 5 µm. Abbreviations: Ctx, cholera toxin subunit B; DAPI, 4’,6-diamidino-2- phenylindole, SIM, structured illumination microscopy; SYP, synaptophysin; WGA, wheat germ agglutinin.

## Discussion

We have investigated the NCU in healthy human skin and provide evidence for crucial morphological phenomena, namely nerve fiber ensheathment by keratinocytes and Cx43 contact sites between keratinocytes and IENF. These findings may profoundly change the view on the role neuronal and non-neuronal cells play in the development and maintenance of neuropathy and neuropathic pain.

Ensheathment was previously described in model organisms with mono- or double-layered epidermis such as *Drosophila* and *Danio rerio* (Han *et al*., 2012; O’Brien *et al*., 2012). In multi-layered mammalian and human skin, only sparse data exist from early EM studies reporting conflicting observations. Given the small diameter of IENF (≤1 µm) within the dense tissue of the epidermis, application of super-resolution microscopy techniques is inevitably necessary to resolve the exact course of these neurites. Recently, “tunneling” of IENF within keratinocytes was proposed via confocal microscopy in human skin (Talagas *et al*., 2020a) and via SEM in an heterologous rat-human co-culture model (Talagas *et al*., 2020b). srAT and ExM techniques used in our study fortify these findings at ultrastructural level and ExM opens the avenue for detailed assessment in diagnostically relevant tissue sections. We further identified Cx43 plaques of keratinocytes in close proximity to IENF as potential components of the NCU exactly size-matching similar connexin and innexin plaques (Agullo-Pascual *et al*., 2013; Markert *et al*., 2016; Taki *et al*., 2018). Keratinocyte-keratinocyte communication via calcium wave propagation and ATP release are canonical functions of Cx43, orchestrating proliferation, wound healing, and inflammatory processes (Martin *et al*., 2014; Tsutsumi *et al*., 2009). It is of note that ATP was also found to be a direct signal transducer from keratinocytes to sensory neurites (Cook and McCleskey, 2002; Sondersorg *et al*., 2014).

Interactions at the NCU may have unprecedented implications for a wide range of somatosensory functions in health and disease. In *Drosophila* larvae, a bidirectional guidance mechanism stabilizing existing fibers and limiting fiber arborization was proposed maintaining sensory receptive fields. In this model, disturbance of ensheathment reduced nocifensive behavior (Jiang *et al*., 2019). In human patients, small fiber pathology is characterized by functional and/or morphological impairment of IENF and is a common finding in a range of neurodegenerative, metabolic, and chronic pain-associated diseases (Ghasemi and Rajabally, 2020; Pittenger *et al*., 2005; Vinik *et al*., 2001). Skin punch biopsies are an easily accessible biomaterial of increasingly acknowledged diagnostic value (Evdokimov *et al*., 2019; Lin *et al*., 2016). Hence, studying the NCU may help understand the pathophysiology of diseases of the peripheral and central nervous system including dystrophic changes typically found in skin of patients with neuropathies (Hovaguimian and Gibbons, 2011).

It is pivotal to recognize and further investigate the active role of keratinocytes within the NCU. Keratinocytes communicate with IENF via ATP and facilitate normal and nociceptive sensory perception in mechanical and thermal modalities (Moehring *et al*., 2018; Sadler *et al*., 2020). Vesicles (Maruyama *et al*., 2018), pannexins (Sondersorg *et al*., 2014), and connexins (Barr *et al*., 2013) are potential mediators of this ATP release and might be dependent on the evoking stimulus. Our observation of Cx43 plaques along the course of IENF in native skin and human co-culture model substantiates a morphological basis for keratinocyte hemichannels as signaling pathway towards IENF. A single cell RNA-sequencing approach of human epidermal cells determined “channel keratinocytes” with upregulated pore and intercellular communication transcripts, e.g. Cx26 and Cx30 (Cheng *et al*., 2018). Hemichannel or even gap junctional communication between keratinocytes and IENF might hence not be restricted to Cx43 and differentially organized in varying specialized keratinocytes.

We successfully established a fully human co-culture model of sensory neurons and keratinocytes maintaining viability for at least six days and neurites growing towards and interacting with keratinocytes. The 2D culture system reduced the multilayered complexity of native skin, yet conserved ensheathment and Cx43 plaques as hallmarks of the NCU. Embryonic stem cell-derived human sensory neurons and human keratinocytes were successfully co-cultured before resulting in direct contacts and engulfed neurites (Krishnan-Kutty *et al*., 2017). However, our model uses fibroblast-derived iPSC which can be generated from virtually any relevant group of patients. Recently, a heterologous model of rat DRG neurons and human keratinocytes showed neurites passing along a keratinocyte gutter or being ensheathed by keratinocytes, which matches our observations. Additionally, SYP, synaptotagmin, and syntaxin 1A were successfully labeled demonstrating synapse-like contacts together with cytokeratin 6 as a keratinocyte marker and pan-neurofilament as a neurite marker (Talagas *et al*., 2020b). Neurofilaments represent intermediate filaments within the cytoplasm and may not encompass the whole neuronal outline compared to membrane labeling (see Figure supplement 3). In accordance with findings from native DRG neurons, our sensory iPSC neurons contained SYP accumulations within the cytoplasm (Chou *et al*., 2002; Chung *et al*., 2019), deeming a membranous labeling necessary to clearly localize SYP in a co-culture approach. Keratinocytes showed no clusters of SYP associated to passing neurites (Figure 8). Still, electrophysiological activity of neurons in contact with keratinocytes can be attenuated by blocking vesicular secretion of keratinocytes via botulinum neurotoxin type C, hinting towards a physiological role in signal transduction at the NCU (Talagas *et al*., 2020b).

Our data further show that diagnostic interpretation of IENF density in skin punch biopsies solely based on PGP9.5 labeling deserves some caution. PGP9.5 is a cytoplasmic marker that may not be distributed homogenously along the whole fiber and omits the neurites membrane (Figure 1), which will be important for localization and co-localization of involved proteins at super-resolution. Regarding *in vitro* approaches, our homologous model should minimize variability and potential artifacts of heterologous co-cultures, especially since profound differences in neuronal DRG subpopulations can be observed across mammalian species (Klein *et al*., 2021; Kupari *et al*., 2021; Shiers *et al*., 2020).

### Conclusion

Sophisticated cell culture and animal models along with super-resolution microscopy are barely beginning to unveil the complexity of the NCU. Epidermal keratinocytes show an astonishing set of interactions with sensory IENF including ensheathment and electrical and chemical synapse-like contacts to nerve fibers (Figure 9). Our morphological findings underline the significance of keratinocytes in somatosensorics and cutaneous nociception and add to the increasing change of textbook knowledge viewing sensory fibers as the sole transducers of environmental stimuli. Expanding investigations towards skin cell impairment in small fiber pathology will help to better understand the underlying mechanisms and open new avenues for targeted treatment.

**Figure 9.**
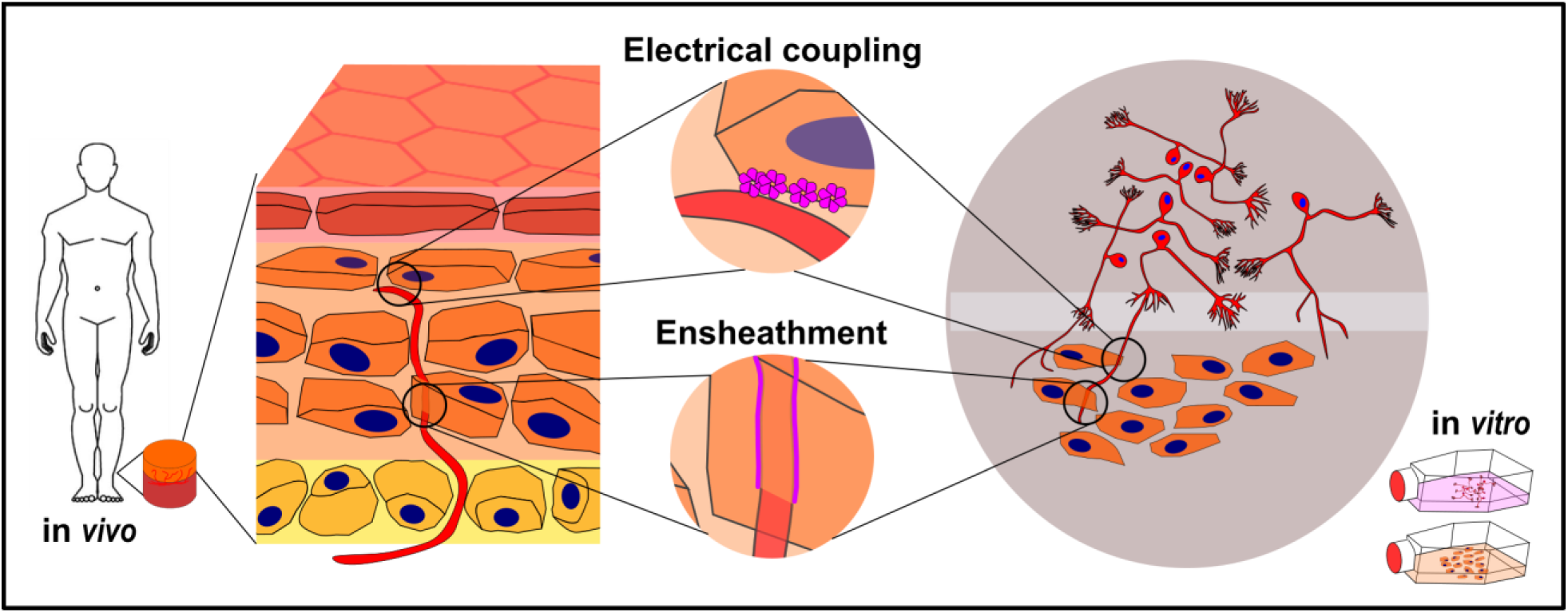
Keratinocyte-nerve fiber interactions in human epidermis and 2D model. Proposed electrical synapses are potential transducers of sensory and nociceptive keratinocyte adenosine triphosphate signaling towards intraepidermal nerve fibers. Ensheathment of fibers by keratinocytes may orchestrate nerve fiber outgrowth and stabilization. Both observations are conserved in human 2D co-culture model.

## Acknowledgements

We thank Dr. Franziska Karl-Schöller for technical help in co-culture handling, Viktoria Diesendorf, BSc for routine sensory neuron differentiation and Alexandra Gentschev, BSc for srAT test runs (all Department of Neurology, University of Würzburg, Germany).

We also thank Daniela Bunsen, Claudia Gehrig-Höhn (Biocenter, Imaging Core Facility, University of Würzburg, Germany), and Dr. Sebastian Markert (Department of Cell Biology, Johns Hopkins University, Baltimore, USA) for expert technical help in srAT. We further thank Dr. Jan Schlegel (Department of Biotechnology and Biophysics, University of Würzburg, Germany) for the preparation of SeTau647 conjugated secondary antibody and Dr. Ralph Götz (Department of Biotechnology and Biophysics, University of Würzburg, Germany) for the introduction into expansion microscopy. The study was supported by the German Research Foundation (Deutsche Forschungsgemeinschaft, DFG UE171/4-1). N.Ü. was supported by DFG UE171/15-1. M.S. was supported by the grant ULTRARESOLUTION from the European Research Council.

## Author contributions

C.E, N.Ü, and C.S. conceptualized the projects and experiments. C.E. performed all experiments. T.K. established methodology for stem cell generation and neuronal differentiation. S.B. and P.D. supported srAT sample generation and imaging. C.E. and N.Ü. wrote the manuscript with contributions from S.B., M.S., and C.S. All authors read and approved the manuscript.

---

**Figure supplement 1:**
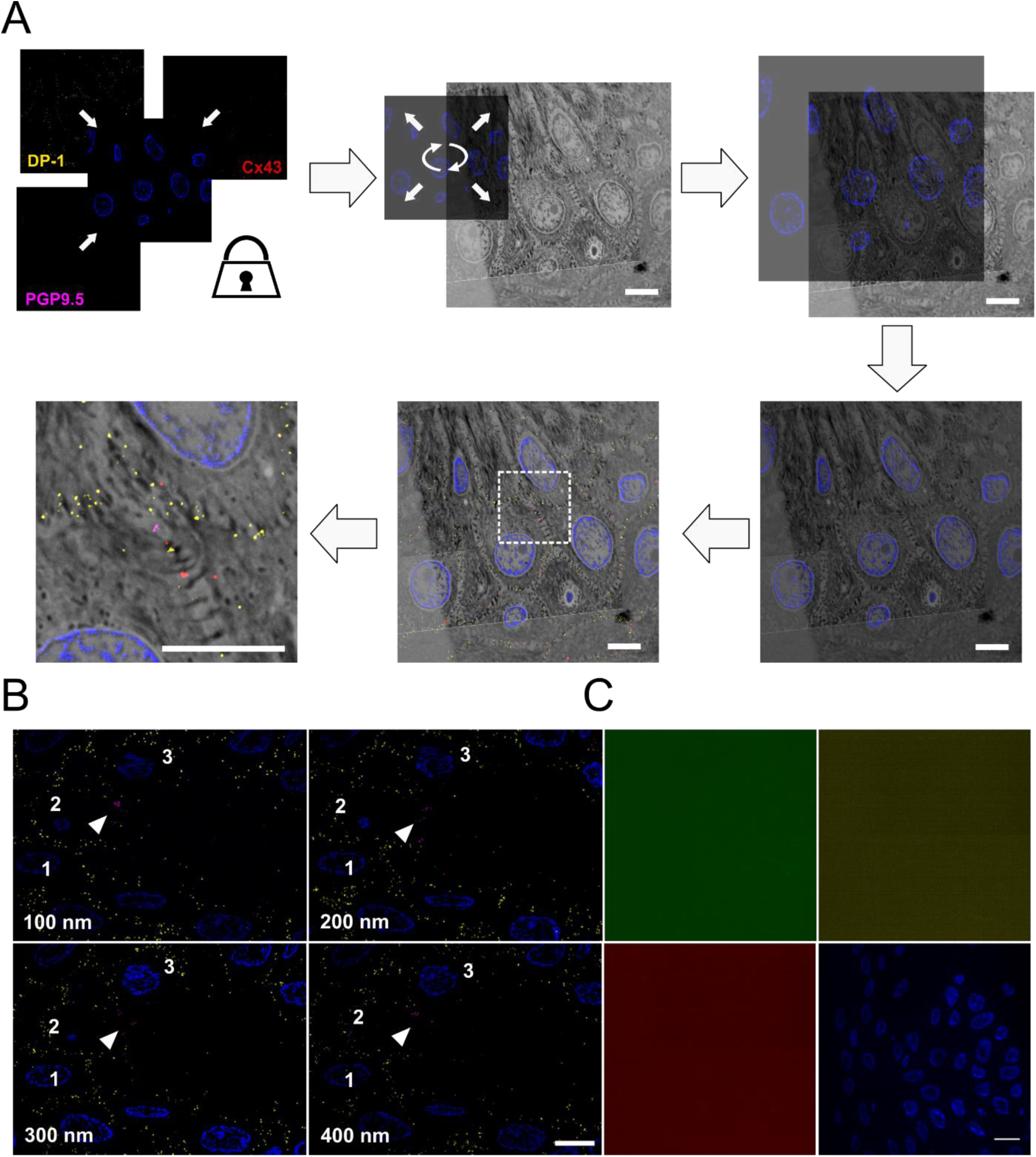
CLEM principle and labeling verification. **(A**) Principle of unbiased correlation using intrinsic landmarks. IF channels are locked and hidden behind nuclear DAPI signal, which is used as only visible fluorescent channel, applied as landmark channel (blue). DAPI signal and SEM nucleus texture are used to achieve unbiased correlation of IF and SEM information. After optimization of the correlation, the channels of interest are made visible, overlaid and regions of interest selected. No further changes of image locations were applied afterwards. IF signal specificity. (B) Four consecutive tissue sections with nuclear DAPI (blue), DP-1 (yellow), and PGP9.5 (magenta) labeling showing traceable persistent labeling. Arrowhead indicates PGP9.5 positive nerve fiber and numbers show position of respective nuclei. Nucleus number 2 is ceasing within the z-stack. Scale bar 5 µm. (C) Application of secondary antibodies Al488-anti mouse (green), SeTau-647-anti rabbit (yellow), and Cy3-anti guinea pig (red) alone showed no fluorescent signal. DAPI signal (blue) for orientation. Scale bar 10 µm. Abbreviations: Al488, Alexa Fluor 488; Cy3, cyanine 3; DAPI, 4’,6-diamidino-2-phenylindole ; DP-1, desmoplakin 1; IF, immunofluorescence, PGP9.5, protein product 9.5 SEM, scanning electron microscopy.

**Figure supplement 2.**
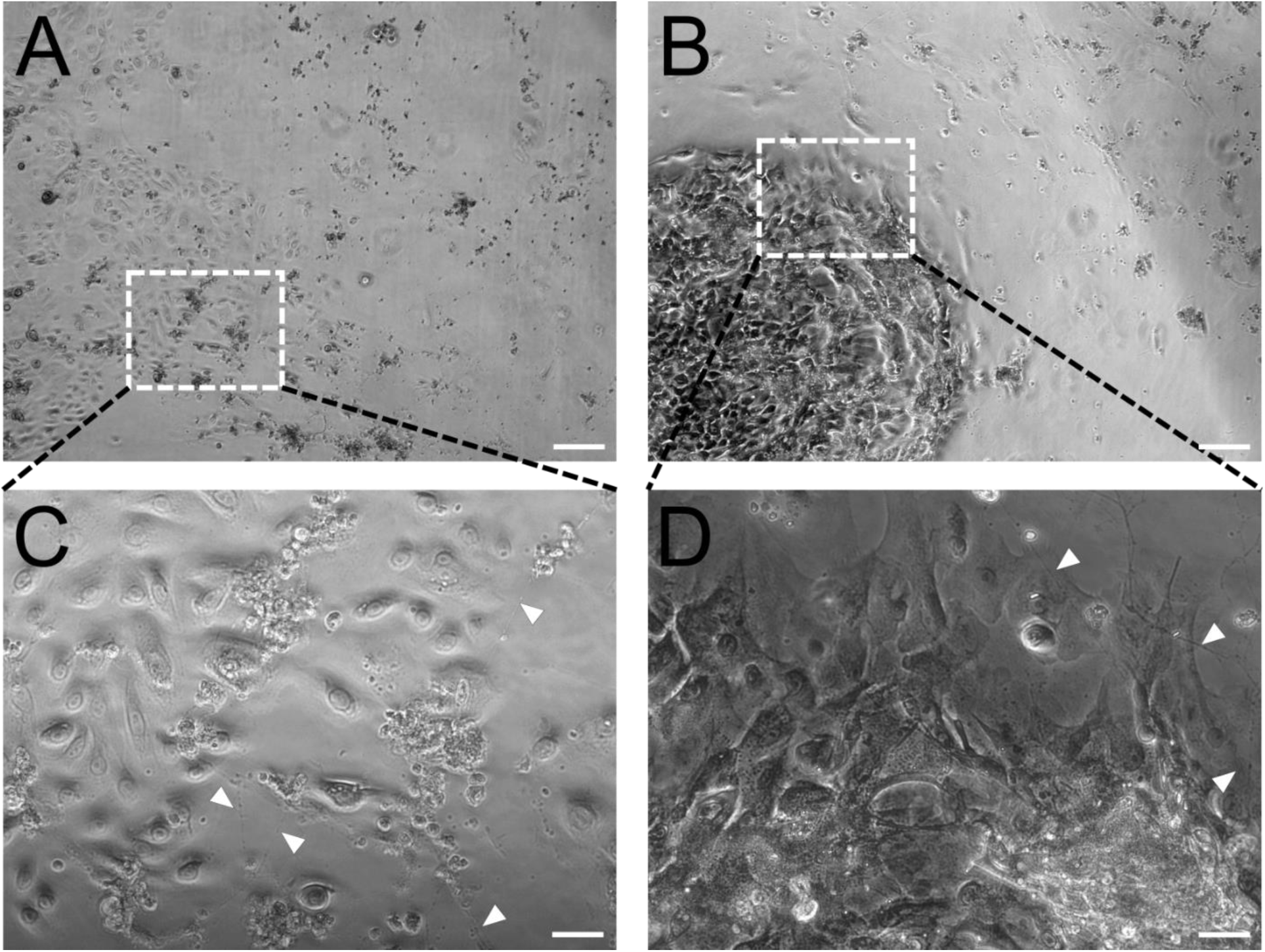
Comparison of co-culture dependent on media condition. Overview of contrast imaged co-culture in keratinocyte medium (A) and conditioned neuronal medium (B) imaged with 5x objective. Insets represent enlarged areas in C and D at 20x objective. Single keratinocytes show basal or early differentiated state in keratinocyte medium (C), whereas differentiated and aggregated keratinocytes are predominant in conditioned neuronal medium (D) Arrowheads indicate passing neurites. Scale bars: 200 µm (A, B) and 50 µm (C, D).

**Figure supplement 3.**
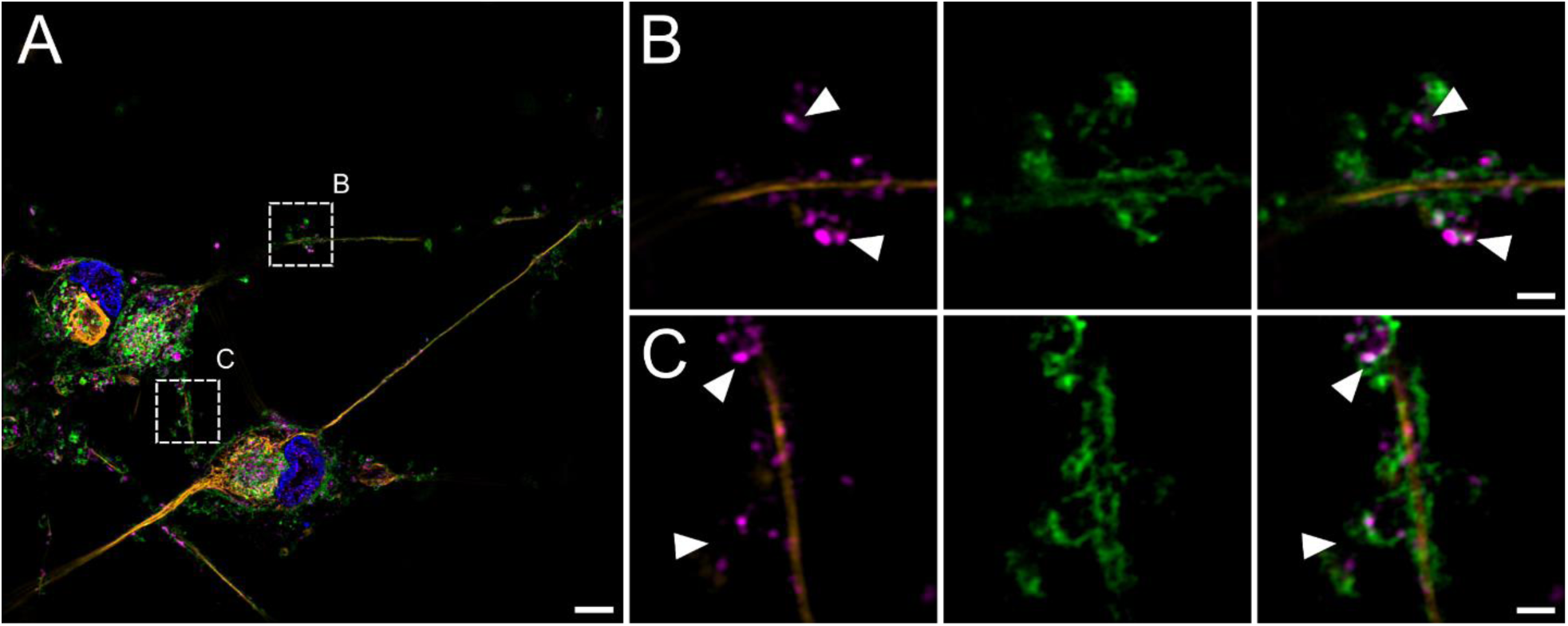
Neurite outline and synaptic vesicular SYP localization. (A) Overview images of neuronal culture with DAPI (blue), Ctx (green), pan-NF (yellow), and SYP (magenta). Insets indicate enlarged areas in B and C. (B, C). First panel shows overlay of neurofilament marker pan-NF and SYP with apparent extra neuronal SYP accumulations (indicated via arrowheads). Second panel shows neurite membrane marker Ctx. Last panel depicts overlay, revealing SYP signal located within the outline of neurites. Scale bars: 5 µm (A), 1 µm (B, C). Abbreviations: Ctx, cholera toxin subunit B; DAPI, 4’,6-diamidino-2-phenylindole, pan-NF, pan-neurofilament; SIM, structured illumination microscopy; SYP, synaptophysin.

## Supplementary Information

**Video 1:** srAT with serial 100 nm sections with two frames per second. PGP9.5 (magenta) marks IENF ensheated in keratinocytes. Scale bar 5 µm. Abbreviations: IENF, intraepidermal nerve fiber; PGP9.5, protein product 9.5; srAT super-resolution array tomography.

**Video 2**. srAT with serial 100 nm SEM sections and 3D interpolation of keratinocyte nuclei (blue), keratinocyte cell bodies (yellow-orange), and IENF (magenta). Four sections per second. Scale bar 5 µm. Abbreviations: IENF, intraepidermal nerve fiber; SEM, scanning electron microscopy; srAT super-resolution array tomography.

**Video 3**. Ensheathment and Cx43 plaque in expanded human epidermis. Magnification with filled arrowhead indicating part of an IENF, labeled via PGP9.5 (magenta), tunneling through a keratinocyte, labeled via phalloidin (grey). Additional hollow arrowhead showing a Cx43 plaque in contact with IENF. Physical 1.2 µm stack step size, translating to 276 nm biological step size. Framerate of two planes per second. Expansion factor corrected scale bar 1µm. Abbreviations: Cx43, connexin 43; IENF, intraepidermal nerve fiber; PGP9.5, protein gene product-9.5.

**Video 4**. Live imaging of fully human sensory neuron-keratinocyte co-culture with neurite establishing contact to keratinocyte colony. Arrowheads indicate outgrowing neurite. Framerate of eight time points per second with 20 min intervals per time point. Scale bar 50 µm.

